# CXCL13/CXCR5 Chemokine Axis Promotes CXCR5^+^CD19^+^ B-Cell and Follicular/Effector CXCR5^+^CD4^+^ T-Cell Responses in the Lungs Associated with Protection from Severe and Fatal COVID-19 Following Infection with Pathogenic SARS-CoV-2 Delta Variant

**DOI:** 10.1101/2025.04.18.649623

**Authors:** Hawa Vahed, Aziz Chentoufi, Swayam Prakash, Afshana Quadiri, Sweta Karan, Yassir Lekhbach, Etinosa Omorogieva, Swena Patel, Jimmy Tadros, Emma Jane Liao, Lauren Lau, Delia F Tifrea, Lbachir BenMohamed

**Author notes:** Authors have contributed equally to this study (Aziz Chentoufi, Swayam Prakash). Corresponding author: Laboratory of Cellular and Molecular Immunology, Gavin Herbert Eye Institute; Hewitt Hall, Room 2032; 843 Health Sciences Rd; Irvine, CA 92697-4390. <u>Support and Conflict of Interest:</u> Studies of this report were supported by Public Health Service Research grants AI158060, AI150091, AI143348, AI147499, AI143326, AI138764, AI124911, and AI110902 from the National Institute of Allergy and Infectious Diseases (NIAID) to LBM. LBM has an equity interest in TechImmune, LLC., a company that may potentially benefit from the research results and serves on the company’s Scientific Advisory Board. LBM’s relationship with TechImmune, LLC., has been reviewed and approved by the University of California, Irvine, under its conflict-of-interest policies.

## Abstract

Chemokines play an important role in shaping lung innate and adaptive immunity to pulmonary infections and diseases. However, the role of CXC ligand 13 (CXCL13), a chemokine homeostatically produced by various lung cell types, in the protection from SARS-CoV-2 infection and disease remains controversial. Some studies reported that asymptomatic patients who survived severe COVID-19 had CXCL13-dominated mucosal immune responses in the lungs early during infection. In contrast, other studies reported that a high level of CXCL13 was associated with severity and mortality in COVID-19 patients. In this study, to determine the direct role of CXCL13 in SARS-CoV-2 infection and disease, we generated CXCL13***^-/-^***K18-hACE2 mice, that are both transgenic for ACE2 and deficient in CXCL13 and compared their infection and COVID-19-like disease symptoms with those in wild-type K18-hACE2 transgenic mouse littermates following intranasal inoculation with the pathogenic SARS-CoV-2 delta variant (B.1.617.2). Compared to age- and gender-matched SARS-CoV-2 infected wild-type K18-hACE2 mice, SARS-CoV-2 infected CXCL13***^-/-^***K18-hACE2 deficient mice exhibited (*i*) higher viral load in the lungs; (*ii*) severe COVID-19-like lung pathology; (*iii*) exacerbated weight loss; (*iv*) increased mortality. The apparent severe COVID-19-like symptoms in CXCL13^-/-^K18-hACE2 deficient mice were associated with: (*i*) significantly lower frequencies of functional lung-resident C-X-C chemokine receptor 5^+^ (CXCR5)^+^CD19^+^ B cells, follicular CXCR5^+^CD4^+^ helper T cells (Tfh cells), and IFN-γ^+^TNF-α^+^GzmB^+^Ki67^+^effector CD4^+^ Th_1_ cells; and (*ii*) a significant reduction in the levels of SARS-CoV-2-Spike specific Th1 associated IgG_1_ and IgG_2b_ antibody isotypes. These findings corroborate previous human reports suggesting a critical role of the CXCL13/CXCR5 chemokine axis in the protective B- and T-cell mucosal immunity to SARS-CoV-2 infection and disease, offering a potential new immunotherapeutic target for treatment.

## INTRODUCTION

Severe acute respiratory syndrome coronavirus 2 (SARS-CoV-2) primarily spreads through aerosols when an infected person coughs, sneezes, or talks^1–4^. The infectious virus can also spread via surface contact with the virus, followed by touching the mouth, nose, or eyes ^5^. Since 2019, many variants of concern (VOCs) of SARS-CoV-2 have continuously emerged: Alpha (B.1.1.7), Beta (B.1.351), Gamma (P.1), Delta (B.1.617.2), and Omicron (B.1.1.529) ^6–11^. COVID-19, caused by SARS-CoV-2 infection of the respiratory tract, can manifest within 1-2 weeks of exposure, with some people remaining asymptomatic while others develop shortness of breath to flu-like symptoms such as fever, cough, loss of taste and smell, and pneumonia ^12^. While many infected individuals remain asymptomatic, making virus transmission challenging to control ^13^, in symptomatic individuals, the disease can extend to acute respiratory distress syndrome and even multi-organ failure, leading to high morbidity and mortality ^4^.

Mucosal tissues, including the respiratory tract mucosal surface, are frontline barriers continuously exposed to infectious pathogens, such as SARS-CoV-2 ^14^. Respiratory tract-resident B- and T-cells play a crucial role in the defense against invading mucosal pathogens and constantly traffic into and surveil these mucosal tissues. A currently accepted paradigm is that tissue-selective trafficking of memory and effector B- and T cells to mucosal tissues is controlled by specific combinations of adhesion receptors and chemokines, often in a tissue-specific manner ^15–23^. Chemokines are a family of small secretory proteins, classified into four subfamilies based on CC, CXC, and CX3C, as well as C chemokines, based on conserved N-terminal cysteine residues ^24^. Several clinical investigations have revealed that many chemokine/chemokine axes mediate the recruitment of immune cells to infected lungs and are associated with either severe symptomatic or asymptomatic COVID-19 ^25–30^. The CXC ligand 13 (CXCL13), also known as B cell chemotactic factor (BLC), ^31^ is the homeostatic chemokine produced by various cell types in the lungs, including follicular dendritic cells, stromal cells, fibroblasts, alveolar macrophages, and monocyte-derived macrophages. It may be pivotal for the migration and proliferation of B- and T-cells in the infected lungs ^32^. Along with its receptor CXCR5, CXCL13 facilitates B cell homing and maintenance of B cell follicles in secondary lymphoid tissues, such as the lungs ^33–35^. The CXCL13-CXCR5 axis plays a beneficial role in virus clearance by promoting B cell differentiation in the ectopic germinal center (GC) to generate broadly neutralizing protective antibodies ^36^. Nessrine Bellamri reported in 2020 that the CXCL13 chemokine is produced by alveolar macrophages from idiopathic pulmonary fibrosis (IPF) patients ^37^. However, the role of the CXCL13-CXCR5 axis in the protection or pathogenicity of SARS-CoV-2 infection and disease remains controversial. One study reported that early expression of CXCL13 in ICU and non-ICU individuals may reflect a potent host immune response, promoting the maturation of B cells and antibody production to control viral replication and achieve virus clearance ^38^. Another study suggested that CXCL13 expression early during the infection was associated with the survival of patients with severe COVID-19. In contrast, CXCL13 expression was absent in patients who progressed into severe and fatal disease ^39^. Other reports have shown that high levels of CXCL13 in the sera were associated with severe COVID-19 prognosis and increased mortality following SARS-CoV-2 infections, thus suggesting CXCL13 as a biomarker for predicting disease progression ^40^. Persistently elevated levels of CXCL13 in the tissue and sera could be detrimental by promoting lung inflammation and tissue fibrosis ^36^. Thus, while chemokines play a role in lung immunity and immunopathology to SARS-CoV-2 infection and disease, the role of CXCL13 chemokine remains to be fully elucidated.

In this study, we investigated the direct role of CXCL13 in SARS-CoV-2 infection and disease. For this purpose, we generated CXCL13***^-/-^***K18-hACE2 mice that are both transgenic for ACE2 and deficient in CXCL13 and compared their infection and disease with wild-type K18-hACE2 mice following infection with the pathogenic SARS-CoV-2 delta variant (B.1.617.2). We demonstrated that the absence of CXCL13 chemokine exacerbated SARS-CoV-2 infection and disease associated with significantly lower frequencies of functional lung-resident CD19^+^CXCR5^+^ B cells, CD4^+^CXCR5^+^ follicular helper T cells (Tfh cells) in the infected lungs and a significant reduction of Th1 cell response known for its protective role in SARS-CoV-2 infection.

## MATERIALS AND METHODS

### Viruses

SARS-CoV-2 viruses specific to the Delta (B.1.617.2) (Batch number: G87167) variant were procured from Microbiologics (St. Cloud, MN). The initial batches of viral stocks were propagated to generate high-titer virus stocks. Calu-3 (ATCC, HTB-55) cells were obtained from ATCC and cultured following an earlier published protocol ^41^. All procedures were completed only after obtaining appropriate safety training using an aseptic technique under BSL-3 containment.

### Mice

Mice aged 7-8 weeks were used in the present study. CXCL13 knockout (NTMCK-211203-ACS-01-tac) mice were purchased from the Taconic Biosciences (Germantown, NY), and WT K18-hACE2 mice (JAX:034860) were purchased from the Jackson Laboratory (Bar Harbor, ME, USA). To obtain CXCL13^-/-^K18-hACE2 deficient mice on the JAX034860 genetic background, CXCL13 knockout mice were crossed to WT K18-hACE2 transgenic mice, which were C57BL/6 background mice. The following primers were used to genotype the mutant and WT *CXCL13* genes in the in-house bred CXCL13^-/-^K18-hACE2 deficient mice colony to check for the CXCL13 and hACE2 genes. Respectively: CXCL13 Forward, 5’-TCACACATGGGCTAGAACGG-3’; Reverse, 5’-TATGCAACGGAGCTTGAGCA-3’; and hACE2 Forward, 5 GGGATCAGAGATCGGAAGAAGAAA-3’; Reverse, 5’-AGGAGGTCTGAACATCATCAGTG-3’. The mice were maintained and bred under specific pathogen-free conditions with a 12-hour light/dark cycle and access to food and water *ad libitum*. The studies were performed following the guidelines of the Assessment and Accreditation for Laboratory Animal Care-accredited Biosafety Level (BSL) 2 and 3 facilities at the NIAID/NIH using a protocol approved (IACUC protocol # AUP-22-086) by the NIAID Animal Care and Use Committee.

### Infection of CXCL13^-/-^K18-hACE2 deficient mice and WT K18-hACE2 mice models

The male CXCL13^-/-^K18-hACE2 deficient mice and WT K18-hACE2 mice were intranasally infected with 5 x 10^5^ pfu of SARS-CoV-2 (Delta B.1.617.2) variant, delivered in 20μl sterile PBS. Mice were monitored daily for death and weight loss until day 14 post-infection (p.i..). On days 2, 6, and 14 days p.i., mice were euthanized for virological and immunological studies.

### Immunohistochemistry of mice lungs

The CXCL13^-/-^K18-hACE2 deficient mice and WT K18-hACE2 mice’s lung sections were preserved in 4% PFA for 48 hours and then transferred to 70% ethanol. The tissue sections were then embedded in paraffin blocks and sectioned at 8 μm thickness. Slides were deparaffinized and rehydrated before hematoxylin and eosin (H&E) staining for immunopathological analysis. Images were captured using the BZ-X710 All-in-One fluorescence microscope (Keyence).

### Virus titration in oropharyngeal swabs

Oropharyngeal swabs from CXCL13^-/-^K18-hACE2 deficient mice (*n* =10) and WT K18-hACE2 mice (n=10) after 2, 6 10, and 14 days p.i. were analyzed for SARS-CoV-2 specific RNA by qRT-PCR. As recommended by the CDC, we used *ORF1ab-*specific primers (Forward-5’-CCCTGTGGGTTTTACACTTAA-3’ and Reverse-5’-ACGATTGTGCATCAGCTGA-3’) to detect viral RNA levels in oropharyngeal swabs. Briefly, 5 µl of the total nucleic acid eluate was added to a 20-µl reaction mixture (1x TaqPath 1-Step RT-qPCR Master Mix (Thermo Fisher Scientific), with 0.9 mM each primer. The q RT-PCR was carried out using the ABI StepOnePlus thermocycler (Life Technologies, Grand Island, NY). When the Ct-value was relatively high (35 ≤ Ct < 40), the specimen was retested twice and considered positive if the Ct-value of any retest was less than 35.

### Flow cytometry

Single-cell suspensions were generated from both the lungs of each mouse by collagenase treatment (8mg/ml) for 1 hour. The resulting cell suspensions were then passed through a 70-μm cell strainer to obtain a uniform single-cell suspension, which was subsequently used for FACS staining. The following antibodies were used: anti-mouse CD45 (BV786, clone 30-F11 – BD), CD3 (BUV395, clone 145-2C11 – BioLegend), CD19 (BV786, clone 6D5 – BioLegend), CD4 (FITC, clone GK1.5 – BioLegend), CXCR5 (PerCP, clone L138D7 – BioLegend), and Ki-67 (BV-421, clone Ki-67 – BioLegend). Surface staining was performed by adding mAbs against various cell markers to a total of 1 x 10^6^ cells in phosphate-buffered saline containing 1% FBS and 0.1% sodium azide, followed by incubation for 45 minutes at 4°C. The cells were washed three times with FACS buffer (PBS, 1% FBS, and 0.1% sodium azide) and fixed in PBS containing 2% paraformaldehyde (Sigma-Aldrich, St. Louis, MO). A total of ∼100,000 lymphocyte-gated PBMCs were acquired by Fortessa X20 (Becton Dickinson, Mountain View, CA). For the intracellular staining, after surface staining, the lymphocytes were treated with the Cytofix/Cytoperm (Becton Dickinson, USA), and stained for 45 minutes at 4°C with IFN-γ (PerCP, clone XMG1.2 – BioLegend), TNF-α (APC-Cy7, clone MP6-XT22 – BD), Gzym-B (APC, clone QA16A02 – BioLegend), and CXCL13 (APC, clone DS8CX13 – Invitrogen). The cells were washed three times with FACS buffer and fixed in PBS containing 2% paraformaldehyde (Sigma-Aldrich, St. Louis, MO). A total of ∼100,000 lymphocyte-gated PBMCs were acquired on the Fortessa X20 (Becton Dickinson, Mountain View, CA) and analyzed using FlowJo software (v10.10.0) (Becton Dickinson, Ashland). The unstained and FMO controls were included to ensure proper gating.

### Luminex

Lung tissue lysates were collected at Day 2, 6, 14 post-infection, and the production of the CXCL13 was assayed using multiplex cytokine arrays per the manufacturer’s protocols (MCKP1-110K, Millipore Sigma). Samples were acquired on a Magpix system (Luminex).

### Enzyme-linked immunosorbent assay (ELISA)

Serum antibodies specific for the SARS-CoV-2-Spike-protein, collected from CXCL13^-/-^K18-hACE2 deficient mice (*n* =10) and WT K18-hACE2 mice (*n* =10), were detected by ELISA. Ninety-six-well plates (Dynex Technologies, Chantilly, VA) were coated with 100 ng of Spike-protein (Sino Biologicals, USA) per well and incubated overnight at 4°C. The next day, the plates were washed three times with PBS and blocked with 3% BSA (in 1X PBS) for 2 hours at room temperature (RT). After blocking, the plates were incubated with serially diluted serum samples (1:20 - 1:312,500; 100 μl/well) and left overnight at 4°C. Next day, the plates were washed thrice with PBS containing 0.05% Tween-20, and bound serum antibodies were detected using 100 μl/well of 1:20,000 dilution of HRP-conjugated rabbit anti-mouse IgG (61-6520-Invitrogen), 1:5,000 dilution of HRP-conjugated goat anti-mouse IgG1 (A10551-Invitogen), 1:5,000 dilution of HRP-conjugated goat anti-mouse IgG2a (A10685-Invitrogen), 1:5,000 dilution of HRP-conjugated goat anti-mouse IgG2b (M32407-Invitrogen) respectively, for 1 hour at RT. Finally, 100 μl/well chromogenic substrate TMB (ThermoFisher) was added, and the plates were incubated for 10-20 min in the dark. Finally, 100 μl/well stop solution (Thermo Fisher, US) was added, and optical density was measured at 450 nm. The cut-off for seropositivity was set as the mean value plus three standard deviations (3SD) from control sera (without secondary antibody addition). All ELISA studies were performed at least twice.

### Immunohistochemistry for chemokine detection

The lung sections collected from CXCL13^-/-^K18-hACE2 deficient mice and WT K18-hACE2 mice at days 6 post-SARS-CoV-2 infection and mock infection were preserved in 4% PFA for 48 hours and then transferred to 70% ethanol. The tissue sections were embedded in paraffin blocks and sectioned at 8 mm thickness. At the time of staining, sections were dewaxed and treated with antigen retrieval reagent, and stained with anti-CXCL13 (PA5-47018–Invitrogen), anti-CXCL14 clone N3C3–GeneTex), anti-CXCL17 (510614–Invitrogen), anti-CCL25 (7N28–GeneTex), and anti-CCL28 (ab231557–abcam) primary antibodies or with isotype control, followed by incubation with HRP-conjugated secondary antibodies. The peroxidase activity was revealed using chromogen substrate (AEC substrate) and slides were counterstained with H&E. The images were captured on the BZ-X710 All-in-One fluorescence microscope (Keyence).

### Statistical analysis

Data for each assay were compared by ANOVA and Student’s *t*-test using GraphPad Prism version 10 (La Jolla, CA). The physical estimation data were analyzed with the paired t-test using a non-parametric Gaussian distribution based on the Wilcoxon matched-pairs signed rank test. Differences between the groups were identified by ANOVA and multiple comparison procedures as previously described ^9, 42, 43^. Data are expressed as the mean ± SD. Results were considered statistically significant at a *P* < 0.05.

## RESULTS

### Expression of CXCL13 chemokine is induced early in the lungs following SARS-CoV-2 infection

The CXC ligand 13 (CXCL13) is a chemokine homeostatically produced by various cell types in the lungs. We first compared the kinetics of expression of CXCL13 in the lungs of age- and gender-matched CXCL13^-/-^K18-hACE2 deficient mice and their WT K18-hACE2 mouse littermates following SARS-CoV-2 infection. On days 2, 6, and 14 p.i., CXCL13^-/-^K18-hACE2 and WT K18-hACE2 mice were euthanized, and lung single-cell suspensions were analyzed for the level of expression of CXCL13 in B cells, CD4^+^ T cells, and CD8^+^ T cells. We observed the maximum expression of CXCL13 chemokine in the lungs of WT K18-hACE2 mice on day 2 p.i. in all three subsets of the lymphocytes, which gradually decreased by day 6 p.i. and dropped significantly by day 14 p.i. (**Fig. 1A** and **B**). As expected, no CXCL13 chemokine expression was detected in the lungs of CXCL13***^-/-^***K18-hACE2 deficient mice as confirmed by both real time PCR at the mRNA level (**Fig. 1C**) and Luminex at the protein level (**Fig. 1D**). However, for WT K18-hACE2 mice, we observed higher levels of CXCL13-specific mRNA (**Fig. 1C**) and protein (**Fig. 1D**) expression. We also analyzed whether CXCL13 deficiency affects the expression levels of the four major mucosal chemokines CXCL14, CXCL17, CCL25, and CCL28, which are homeostatically expressed in the lungs. Age- and gender-matched SARS-CoV-2-infected CXCL13^-/-^K18-hACE2 deficient mice and WT K18-hACE2 mice were euthanized at various time points p.i., and lung tissues were preserved in 4% PFA, paraffin-embedded, sliced, and stained with anti-CXCL13, CXCL14, CCL25, and CCL28 antibodies. We observed no significant difference in the expression of mucosal chemokines CXCL14, CCL25, and CCL28 in CXCL13^-/-^K18-hACE2 deficient mice compared to the WT K18-hACE2 mice, as shown in the representative data on day 6 post SARS-CoV-2 infection (**Fig. 2**).

**Figure 1.**
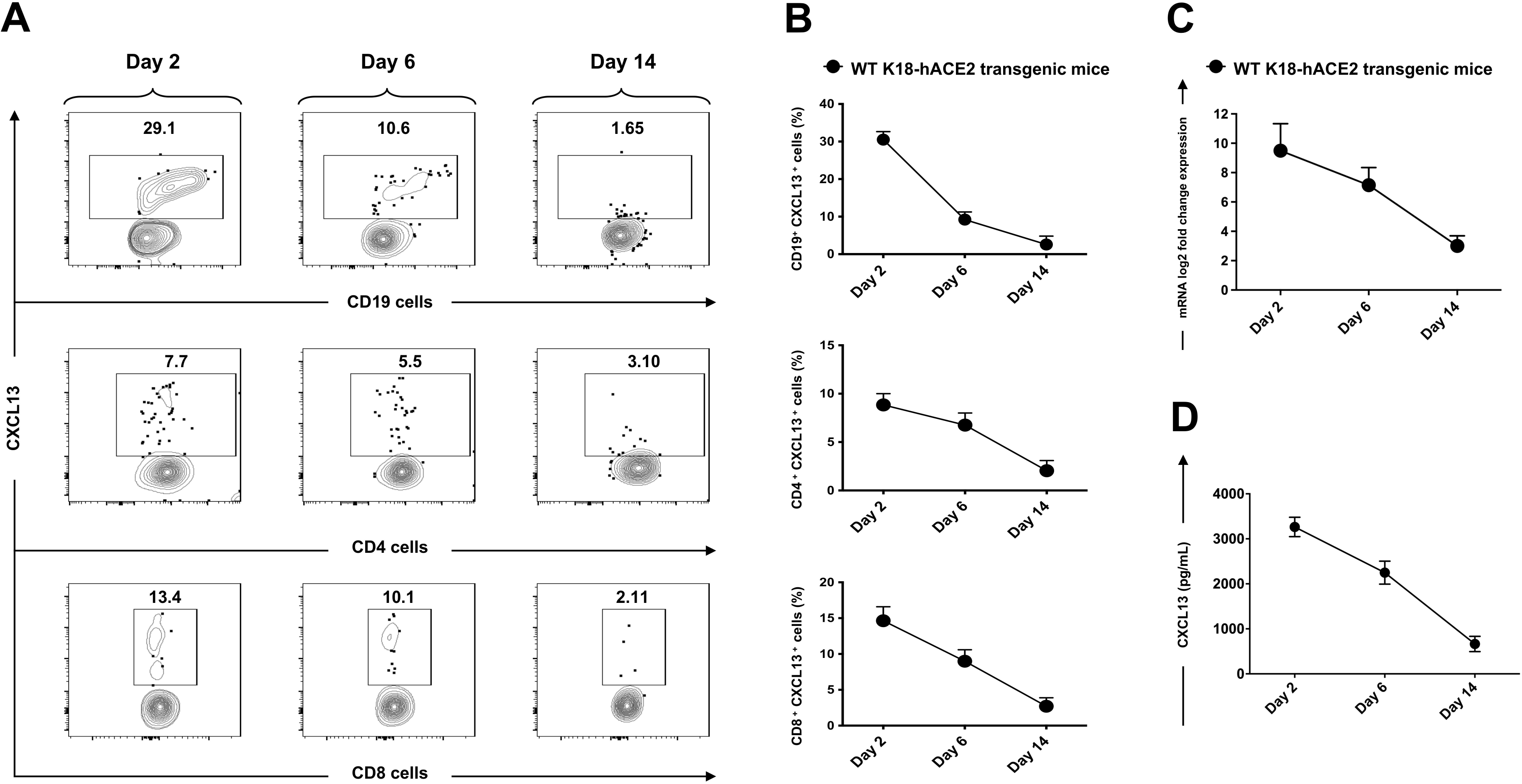
Reduced frequencies and expression of CXCL13 in immune cells and lung tissue in the lungs of WT K18-hACE2 mice following SARS-CoV-2 infection: WT K18-hACE2 mice (*n* = 10) were intranasally infected with 5 x 10^5^ pfu of the SARS-CoV-2 (Delta B.1.617.2) variant. On days 2, 6, and 14 p.i., mice were euthanized, and cell suspension from the lungs was analyzed by flow cytometry. (**A**) Representative FACS plots of frequencies of CD19^+^, CD4^+^, and CD8^+^ cells expressing CXCL13 chemokine in the lungs of WT K18-hACE2 mice after SARS-CoV-2 infection. (**B**) Graphs show differences in the percentage of CD19^+^, CD4^+^, and CD8^+^ cells expressing CXCL13 chemokine in the lung mononuclear cells of mice. (**C**) real time PCR based mRNA expression level, and (**D**) Luminex assay-based protein level of CXCL13 in lung tissue of WT K18-hACE2 mice. Bars represent the mean ± SD.

**Figure 2.**
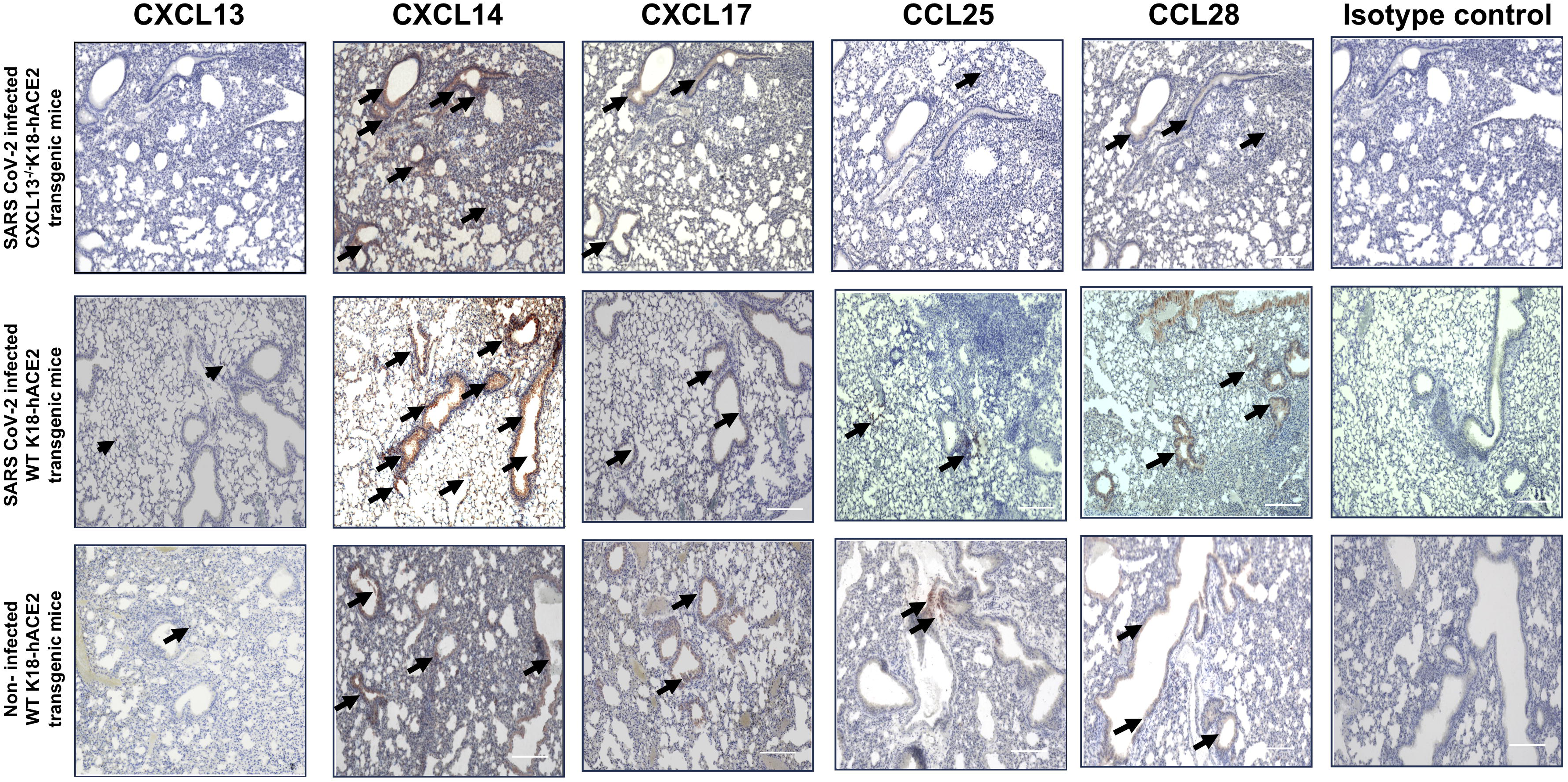
Lung tissue specific expression pattern for mucosal chemokines in SARS-CoV-2 infected CXCL13^-/-^K18-hACE2 and WT K18-hACE2 transgenic mice: Immunostaining of lung sections is shown for SARS-CoV-2-infected CXCL13 deficient CXCL13^-/-^K18-hACE2 transgenic mice (*top panel*), WT K18-hACE2 transgenic mice (*middle panel*) and non-infected WT K18-hACE2 transgenic mice (*bottom panel*). The lung tissue sections were stained with anti-CXCL13, anti-CXCL14, anti-CXCL17, anti-CCL25, and anti-CCL28 primary antibodies or with isotype control, followed by HRP-conjugated secondary antibodies. The peroxidase activity was revealed by chromogen substrate (AEC substrate), and slides were counterstained with H&E. Black arrows point to the positive staining in the lungs showing the chemokine expression. The images were taken at 10X, with a 250μm scale.

### CXCL13 chemokine deficiency is associated with enhanced viral replication and severe COVID-19 pathology in the lungs following infection with the SARS-CoV-2 delta variant

We first determined the functional consequences of CXCL13 deficiency on protection against SARS-CoV-2 infection and disease. For this purpose, we generated CXCL13***^-/-^***K18-hACE2 mice that are both transgenic for ACE2 and deficient in CXCL13 and compared their COVID-19-like disease symptoms with those in wild-type K18-hACE2 mice (**Fig. 3A**). Age- and gender-matched CXCL13^-/-^K18-hACE2 and WT K18-hACE2 mice (*n* = 10) were intranasally inoculated with 5 x 10^5^ pfu of the highly pathogenic Delta variant (B.1.617.2). Compared to WT K18-hACE2 transgenic mice, the CXCL13^-/-^K18-hACE2 deficient mice showed significantly greater weight loss (**Fig. 3B**), and mortality (**Fig 3C).** CXCL13^-/-^K18-hACE2 deficient mice progressively lost up to 12% of their body weight within the first eleven days before gradually returning to their original weight within approximately two weeks (**Fig. 3B**). In contrast, the WT K18-hACE2 mice started losing weight three days post-infection, with a maximum percent body weight loss of 7% observed on day 9 p.i., before gradually returning to their original weight (**Fig. 3B**). Accordingly, by day 14 p.i. with the highly pathogenic Delta variant (B.1.617.2), only 40% of CXCL13^-/-^K18-hACE2 deficient mice survived compared to 80% of WT K18-hACE2 mice (**Fig. 3C**). The severity of Delta variant infection was reflected by significantly higher viral titers detected from the oropharyngeal swabs of the CXCL13^-/-^K18-hACE2 deficient mice compared to WT K18-hACE2 mice (*P* < 0.05). Specifically, there were 1-2 logs higher viral RNA copies in the oropharyngeal swabs of CXCL13^-/-^K18-hACE2 deficient mice compared to WT K18-hACE2 mice detected on days 2, 6, 10, and 14 p.i. with SARS-CoV-2 Delta variant (**Fig. 3D**).

**Figure 3.**
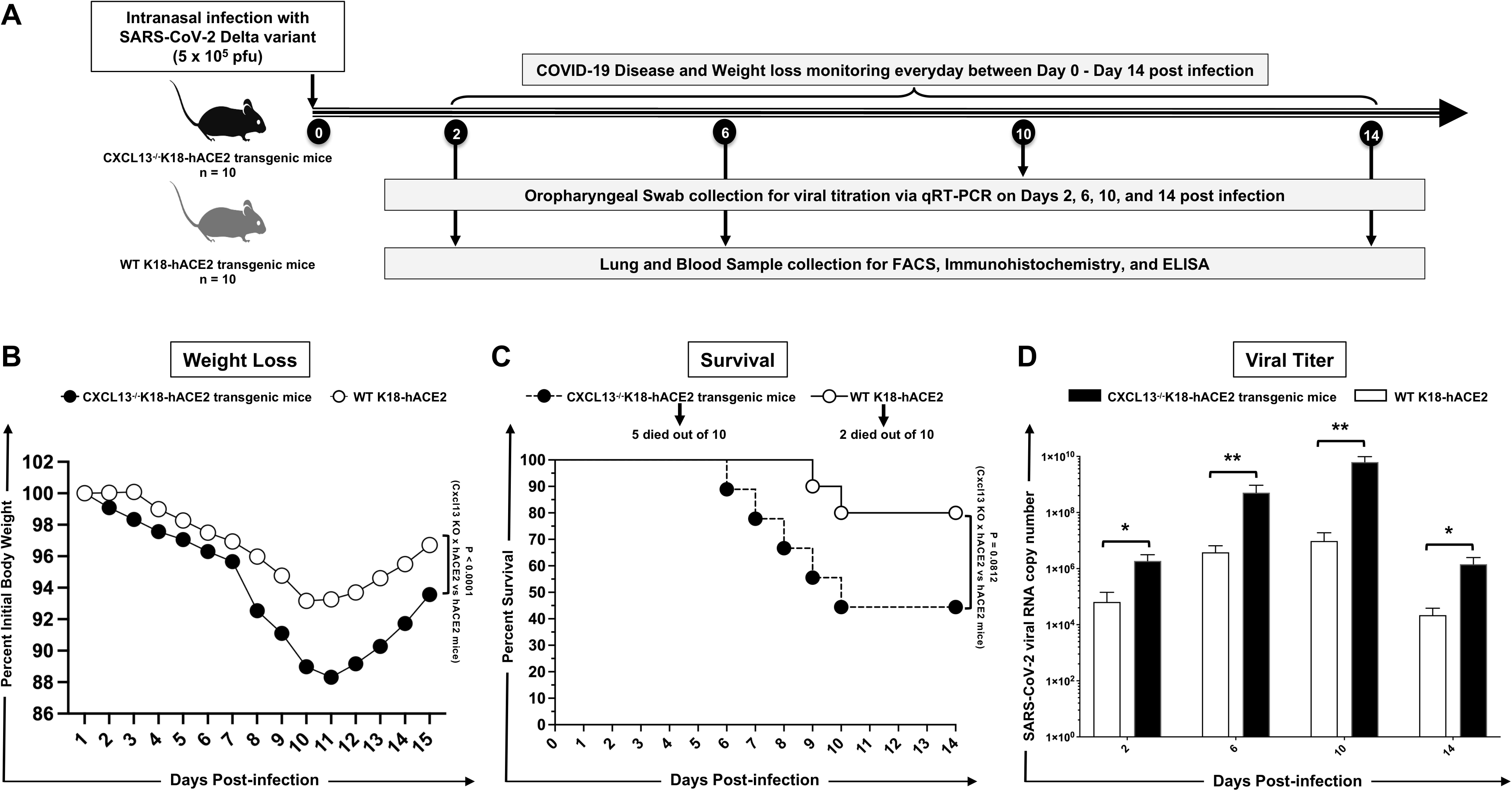
COVID-19 disease and survival in CXCL13^-/-^K18-hACE2 deficient mice and WT K18-hACE2 mice infected with SARS-CoV-2: (**A**) Experimental scheme showing CXCL13^-/-^K18-hACE2 deficient mice and WT K18-hACE2 mice (*n* = 10) were infected intranasally with 5 x 10^5^ pfu of SARS-CoV-2 (Delta B.1.617.2) variant. The weight loss and survival were recorded until 14 days p.i. At days 2, 6, 10, and 14, oropharyngeal swabs were collected for viral copy number through PCR. On day 14 p.i. mice were euthanized, blood was collected, and sera were separated for the ELISA. On days 2, 6, and 14 p.i. mice were euthanized, lungs were harvested, and lung mononuclear cells were isolated for flow cytometry analysis. (**B**) The graph in the left panel shows the average body weight change recorded until 14 days p.i., calculated in percentage, after normalizing to the body weight recorded on the day of infection (day 0). The weight change was monitored until day 14 post-infection. with SARS-CoV-2 infection. (**C**) Shows the percentage survival detected in CXCL13^-/-^K18-hACE2 deficient mice and WT K18-hACE2 mice up to day 14 p.i. (**D**) Data showing viral RNA copy number through PCR in the oropharyngeal swabs at day 2, 6, 10, and 14 days p.i. The data represent two independent experiments; the graphed values and bars represent the SD between the two experiments.

We next determined whether CXCL13 deficiency affects lung pathology. H&E staining of lung sections at days 2, 6, and 14 p.i., with SARS-CoV-2 Delta infection, showed significantly severe COVID-19-related lung pathology in the CXCL13^-/-^K18-hACE2 deficient mice compared to WT K18-hACE2 mice at the mentioned time points. (**Fig. 4)**. Severe lung pathology was observed on Day 6 p.i., in CXCL13^-/-^ K18-hACE2 deficient mice. Still, on Day 14 p.i., hemorrhage, and reduced lung vacuole spaces were observed while studying the lung pathology in CXCL13^-/-^K18-hACE2 deficient mice as compared to WT K18-hACE2 transgenic mice.

**Figure 4.** Lung pathology following SARS-CoV-2 infection in CXCL13^-/-^K18-hACE2 and WT K18-hACE2 transgenic mice: Representative H&E staining images of lungs at day 2, 6, and 14 p.i of SARS-CoV-2-infected mice at 4x and 20x magnifications. Black arrows point to the vacuoles in the lungs. Severe lung pathology was observed in CXCL13 deficient CXCL13^-/-^K18-hACE2 transgenic mice. The images were taken at 4x and 20x, with a 100μm scale.

Altogether, these results indicate that CXCL13 deficiency was associated with higher viral replication in the lungs and severe COVID-19 lung pathology following infection with the pathogenic SARS-CoV-2 delta variant.

### Decreased frequencies of CXCR5^+^CD19^+^ B cells and CXCR5^+^CD4^+^ follicular T helper cells in the lungs of CXCL13^-/-^K18-hACE2 deficient mice following SARS-CoV-2 infection.

We next examined whether CXCL13 deficiency affected the frequencies in the lungs of B- and T cells expressing the C-X-C chemokine receptor 5 (CXCR5), the receptor of CXCL13. On days 2, 6, and 14 p.i., mice were euthanized and cell suspensions from the lung were analyzed by flow cytometry to assess the frequencies of B- and T-cells. We discovered significantly lower frequencies of (*i*) CD19^+^ B-cells (*P* < 0.05) on day 6 p.i. (**Fig. 5A**), (*ii*) CXCR5^+^ CD19^+^ B cells (*P* < 0.05) on day 2 p.i. (**Fig. 5B**), and (*iii*) and CXCR5^+^ CD4^+^ T follicular helper (Tfh) cells (*P* < 0.05) on day 2 and 6 p.i., (**Fig. 5C**) in lungs of CXCL13^-/-^K18-hACE2 deficient mice compared to the frequencies detected in the lungs of age- and gender-matched WT K18-hACE2 transgenic mouse littermates, following infection with SARS-CoV-2 delta variant. There was no significant difference in the frequencies of CXCR5^+^CD8^+^ T cells (*not shown*).

**Figure 5.**
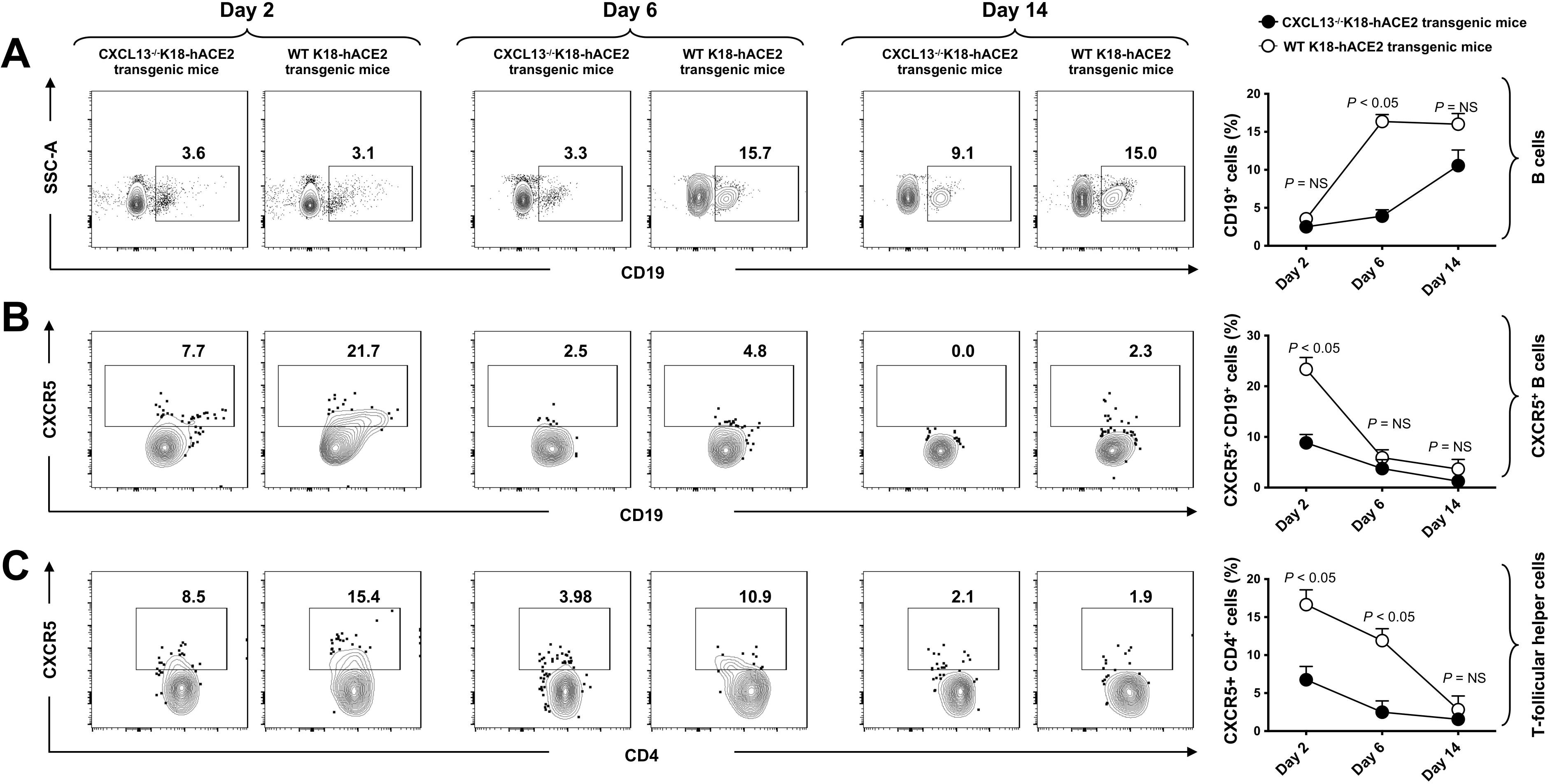
Frequencies of CD19^+^ cells, CD19^+^ CXCR5^+^ cells, and CD4^+^ CXCR5^+^ Tfh cells in the lungs of CXCL13^-/-^K18-hACE2 and WT K18-hACE2 transgenic mice after SARS-CoV-2 infection: Representative and average frequencies of total (**A**) CD19^+^ B cells; (**B**) CD19^+^ CXCR5^+^ cells; and (**C**) CD4^+^ CXCR5^+^ (Tfh) cells in the lungs of CXCL13^-/-^K18-hACE2 mice compared to WT K18-hACE2 mice after SARS-CoV-2 infection. The results shown were representative of two independent experiments. The indicated *P* values were calculated using the unpaired *t*-test.

Altogether, these results suggest that the CXCL13/CXCR5 chemokine axis plays a major role in the mobilization and B cells and CD4^+^ Tfh cells in the lungs to clear SARS-CoV-2 infection and curb COVID-like symptoms, with a yet-to-be-determined mechanism, as illustrated in **Fig. 7**.

### Lack of protection against SARS-CoV-2 infection in CXCL13^-/-^K18-hACE2 deficient mice is associated with reduced frequencies of functional CD4^+^ T helper cells in the lungs.

We next compared the functional marker of CD4^+^ and CD8^+^ T cells in the lungs of the CXCL13^-/-^K18-hACE2 and WT K18-hACE2 mice after SARS-CoV-2 variant infection. On day 14 p.i., mice were euthanized, and lung mononuclear cells were analyzed by flow cytometry to study the expression of IFN-γ, TNF-α, Gzym-B and Ki-67 (cell proliferation marker) in the lung-resident CD4^+^ T helper cells (**Fig. 6A**). However, there was no significant difference observed in the frequencies of CD4^+^ T helper cells (data not shown). Further, we observed significantly reduced frequencies of the IFN-γ (*P* < 0.05), and Gzym-B (*P* < 0.05) producing CD4^+^ T helper cells in the CXCL13^-/-^K18-hACE2 deficient mice compared to WT K18-hACE2 mice (**Fig. 6A**). We also observed significantly lower proliferative responses of Ki-67^+^ CD 4^+^ T helper cells in the lungs of the CXCL13^-/-^K18-hACE2 deficient mice compared to WT K18-hACE2 transgenic mice. This confirmed the role of functional CD4^+^ T helper cells in protection against SARS-CoV-2 infection (*P* < 0.05). Together, these results indicate that during the SARS-CoV-2 infection, CXCL13 induces more IFN-γ and Gzm-B-producing CD4^+^ T helper cells. This conferred more protection against SARS-CoV-2 infection.

**Figure 6.**
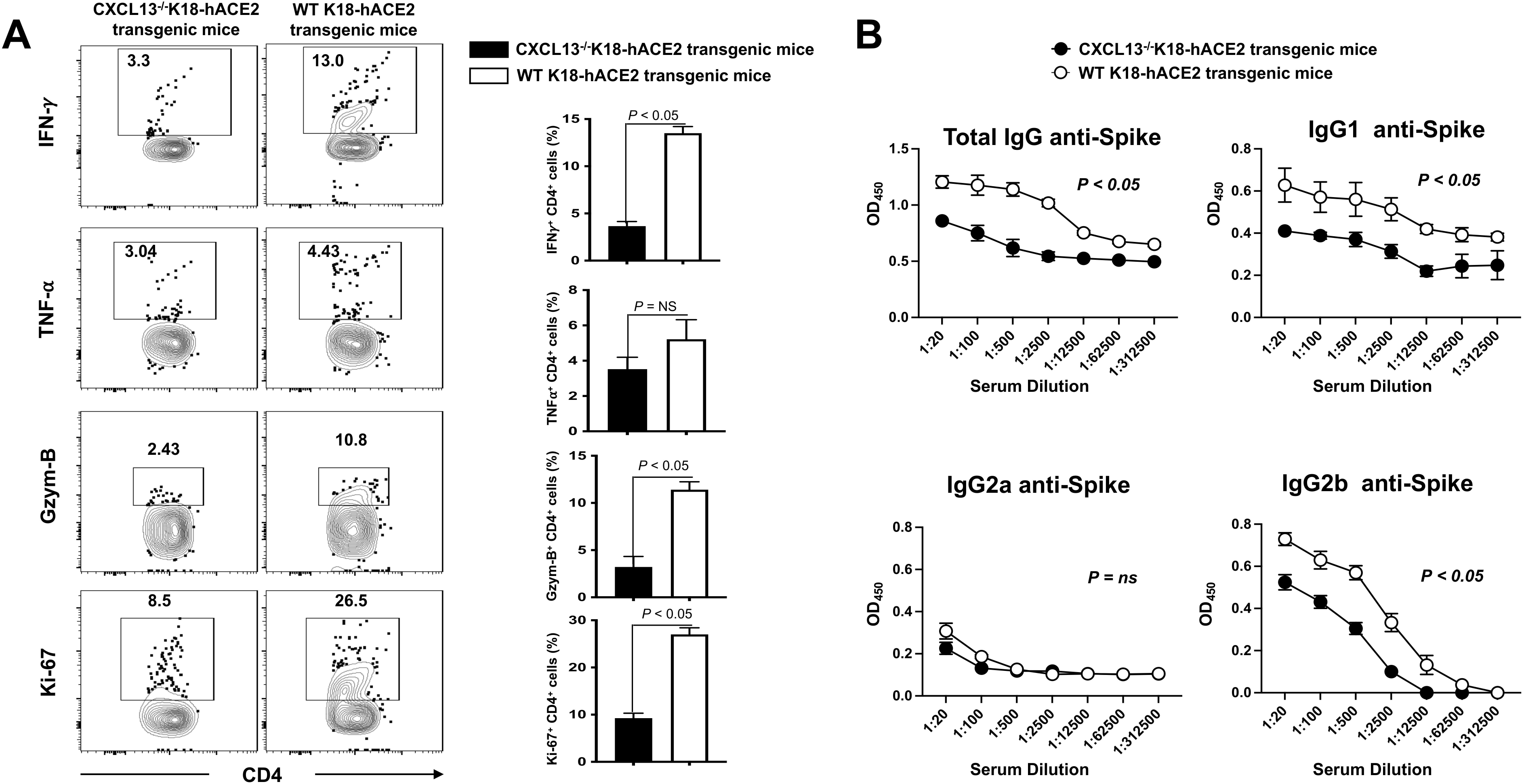
T cell response measured by Cytokines and IgG isotype in the lungs of CXCL13^-/-^K18-hACE2 and WT K18-hACE2 transgenic mice after SARS-CoV-2 infection: (**A**) FACS plots (*left panel*) and Bar graphs (*right panel*) shown for CD4^+^ T cells expressing IFN-γ, TNF-α, Gzm-B, and Ki-67 in the lungs of CXCL13^-/-^K18-hACE2 deficient mice compared to WT K18-hACE2 transgenic mice. Bars represent the mean ± SD. (**B**) Sera from CXCL13^-/-^K18-hACE2 transgenic mice compared to WT K18-hACE2 transgenic mice were analyzed for total IgG, IgG1, IgG2a, and IgG2b response against SARS-CoV-2 (S1+S2 Protein). The upper left graph displays the total IgG levels against SARS-CoV-2 (S1+S2 Protein). The results were representative of two independent experiments. The indicated *P* values were calculated using the unpaired *t*-test.

**Figure 7.**
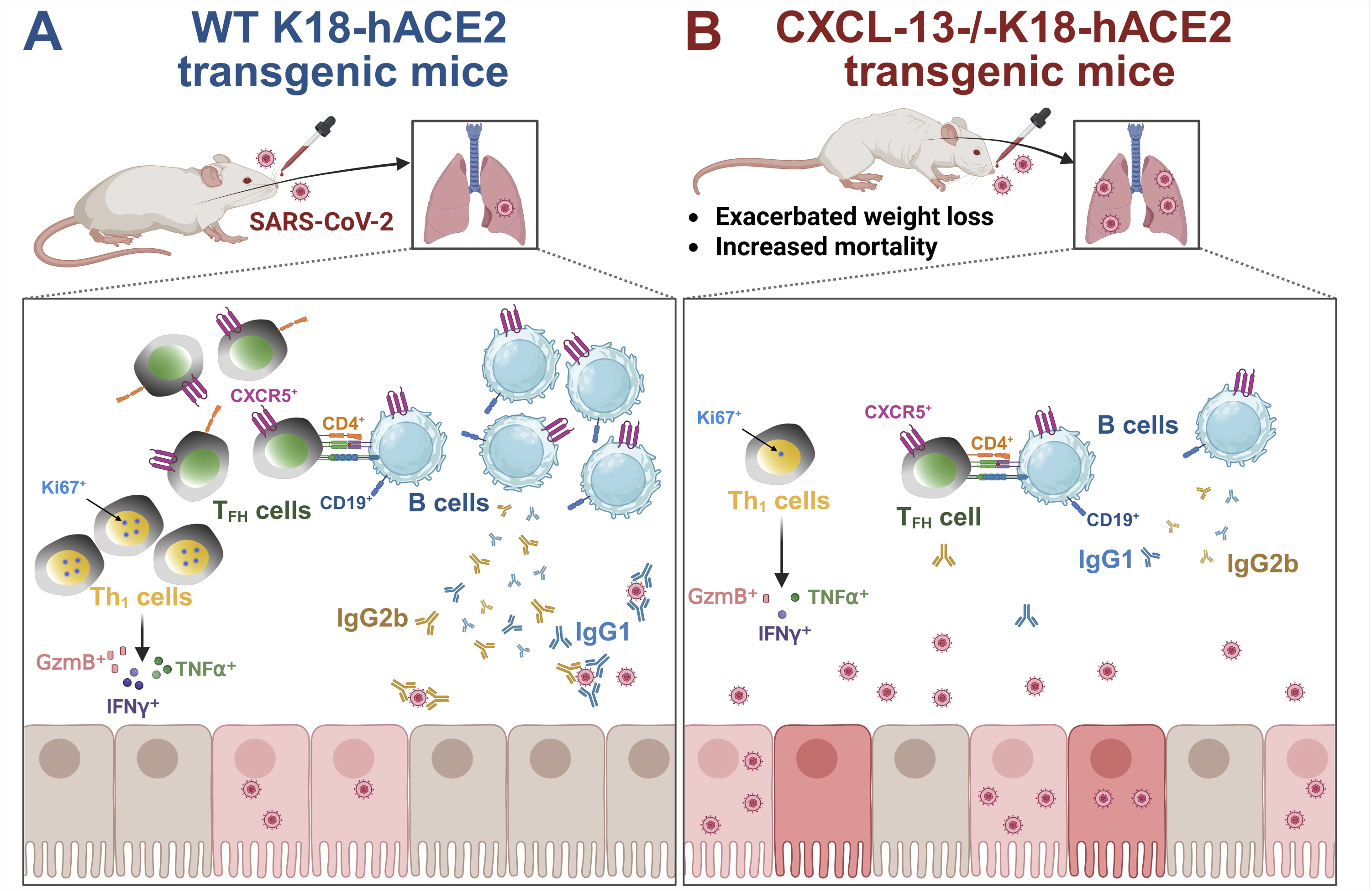
Comparison of immune responses in SARS-CoV-2-infected WT K18-hACE2 and CXCL13^⁻/⁻^ K18-hACE2 transgenic mice: WT hACE2 transgenic mice (*left*) maintain moderate weight and lower viral loads, supported by robust mucosal immune responses, including abundant CD19⁺CXCR5⁺ B cells, CD4⁺CXCR5⁺ Tfh cells, strong IgG1/IgG2b production, and proliferative IFNγ⁺TNFα⁺GzmB⁺Ki67⁺ CD4⁺ Th_1_ cells. CXCL13^⁻/⁻^ mice (*right*) experience pronounced weight loss and elevated viral loads, accompanied by weakened mucosal immunity marked by reduced B and Tfh cell populations, diminished antibody levels, and impaired Th_1_ responses.

### Reduced frequencies of functional lung-resident CD4^+^ T helper cells and lower IgG production detected in symptomatic CXCL13^-/-^K18-hACE2 deficient mice following SARS-CoV-2 infection in

Since we observed a decreased in B cell and Tfh cell frequencies in our CXCL13^-/-^K18-hACE2 deficient mice compared to WT K18-hACE2 mice after SARS-CoV-2 Delta infection, we subsequently compared the IgG levels in both mice models. For this purpose, the antibody titer of SARS-CoV-2 spike-specific IgG, IgG1, IgG2a, and IgG2b was determined by ELISA in the sera collected from CXCL13^-/-^K18-hACE2 deficient mice and WT K18-hACE2 mice infected with the Delta variant of SARS-CoV-2. The WT K18-hACE2 mice showed significantly higher levels of total IgG, IgG1, and IgG2b as compared to CXCL13^-/-^K18-hACE2 deficient mice . However, we did not observe a significant level of expression for IgG2a. Altogether, these results indicate that CXCL13 is essential to induce the cross-protective antibodies that facilitate virus clearance and reduce COVID-19-related disease in mice (**Fig. 6B**). In conclusion, these results indicate that CXCL13 plays a crucial role in the induction of total IgG and its sub-types IgG1, IgG2b and is involved with enhanced protection against SARS-CoV-2 infection.

## DISCUSSION

SARS-CoV-2 infection of the lungs activates a mucosal immune response involving both the innate and adaptive host immune systems ^44^. These mucosal responses lead to the secretion of many soluble mediators, including cytokines, growth factors, and chemokines ^40^. Among the 48 different chemokines, our study focuses on CXCL13 chemokine, a chemokine produced by various cell types in the lungs, including follicular dendritic cells, stromal cells, fibroblasts, alveolar macrophages, and monocyte-derived macrophages ^45–47^. We demonstrated that CXCL13 deficiency exacerbated SARS-CoV-2 infection and disease, leading to severe weight loss, higher mortality, increased viral loads, and severely damaged lungs in CXCL13^-/-^K18-hACE2 deficient mice compared to WT K18-hACE2 mice. Moreover, CXCL13 deficiency was associated with a significant reduction of CXCR5^+^B cells and CXCR5^+^CD4^+^Tfh cells retained in the lungs after SARS-CoV-2 infection. The CXCR5 receptor binds to the CXCL13 ligand, thus facilitating the trafficking of B and Tfh cells into secondary lymphoid tissues ^33,36^. Further, in this study, we observed significantly lower (*i*) levels of SARS-CoV-2-Spike specific Th1-associated IgG1 and IgG2b isotypes; and (*ii*) lower frequencies of IFN-γ and GzmB-producing CD4^+^ T helper cells in infected CXCL13^-/-^K18-hACE2 deficient mice compared to infected WT K18-hACE2 mice (**Fig. 7**). These findings corroborate previous human reports suggesting a critical role of the CXCL13/CXCR5 chemokine axis in the induction of protective B- and T-cell mucosal immunity to SARS-CoV-2 infection and disease, offering a new immunotherapeutic target for treatment.

Recent studies have shown that the elevated levels of the circulating CXCL13 in sera correlate with severe prognosis and increased mortality following SARS-CoV-2 infection, suggesting CXCL13 as a potential biomarker for predicting disease progression and a therapeutic target ^36, 40, 48^. Contrary to this perception, we demonstrate that the CXCL13/CXCR5 chemokine axis was associated with the induction of protective B- and T cell-mediated immunity to SARS-CoV-2 infection and disease, by reducing viral burden in the lungs and curbing lung damage during the infection with the most pathogenic Delta variant of SARS-CoV-2. Our results align with the earlier findings that CXCL13 enhances viral clearance through B cells mediated production of broadly neutralizing protective antibodies ^49^. These neutralizing antibodies are essential for protective immunity against SARS-CoV-2 infection, likely by neutralizing viral particles and preventing virus entry to lung epithelial cells ^50^. However, hospitalized symptomatic patients with severe COVID-19 symptoms persistently showed elevated levels of CXCL13 in the tissue and sera, which might be responsible for promoting inflammation by facilitating inflammatory cell homing and tissue fibrosis in severe COVID-19 disease. Therefore, our preclinical data in CXCL13^-/-^K18-hACE2 deficient mice model do not support targeting CXCL13 during severe COVID-19 infection.

During respiratory viral infections, the upper respiratory tract (URT) serves as the initial site of viral replication, where preexisting adaptive immunity can prevent infection. Interferons (IFNs) and innate immune effector cells can provide an early defense, followed by adaptive T and B cell responses. SARS-CoV-2 infection has been associated with delayed or prolonged innate response, which in turn affects adaptive immune responses. Whether CXCL13 chemokine is expressed by alveolar macrophages of COVID-19 patients, similar to those of idiopathic pulmonary fibrosis (IPF) patients ^37^, remains to be determined. Nevertheless, a recent study showed that patients with severe COVID-19 disease survived due to early interferon (IFN)-dominated mucosal immune responses (IFN-γ, CXCL10, and CXCL13) in infection. These early mucosal responses were absent in patients who progressed to fatal disease despite having similar SARS-CoV-2 viral load ^39^. Additionally, B cells play an important role in the prevention and recovery from COVID-19 disease. The presence of Tfh cells and B cells is crucial for producing functional neutralizing antibodies against SARS-CoV-2 at the early stages of the disease ^51, 52^. Our results support these findings as CXCL13^-/-^K18-hACE2 deficient mice exhibited increased mortality and reduced IFN-γ production. CXCL13 has been implicated in the recirculation of CXCR5^+^ B cells and CD4^+^ T cells and in maintaining chronic inflammation during SARS-CoV-2 infection (**Fig. 7**).

CXCL13 chemokine has been reported to be produced by various cell types in the lungs, including follicular dendritic cells (FDCs) and stromal cells, T cells, B cells, human alveolar macrophages, lung fibroblasts, and monocyte-derived macrophages ^36^. One study reported that CXCL13 is expressed by alveolar macrophages from IPF patients ^37^. Other studies suggest that lung B cells are a major source of CXCL13 ^53, 54^. In our study, CXCL13 was locally expressed in the lung B cells and T cells during SARS-CoV-2 infection in mice. We also found that CXCL13 deficiency did not impact the expression of other mucosal chemokines such as CXCL14, CXCL17, CCL25, and CCL28 in lung tissues of SARS-CoV-2-infected mice. However, the effect of CXCL13 deficiency on pro-inflammatory cytokine secretion revealed a significant decrease in the concentrations of several key pro-inflammatory cytokines, including IL-21, IL-1β, IL-1α, IL-17, IL-6, and TNF-α ^55^. These findings suggest that CXCL13 plays a crucial role in disease pathogenesis and progression ^55^. Although CXCL13 has been reported to be upregulated in SARS-CoV-2 patients, particularly in severe cases, its upregulation was amplified in lethal disease, identifying CXCL13 as a potential novel biomarker for COVID-19 severity ^48^. CXCL13 deficiency has been shown to restrict the migration of CD4⁺CXCR5⁺ T cells in mesenteric lymph nodes, thereby increasing the proportion of regulatory B cells ^55^. Furthermore, studies using CXCL13 knockout mice in the Experimental Autoimmune Encephalomyelitis (EAE) model demonstrated reduced inflammatory infiltration and demyelination in the central nervous system, as well as decreased gliosis and fibrosis ^56^. Therefore, CXCL13 deficiency could influence the secretion of other chemokines and cytokines, potentially affecting the immune response during SARS-CoV-2 infection.

Our study has several significant limitations, as the CXCL13^-/-^K18-hACE2 transgenic mouse model may have adapted to constitutive and complete CXCL13 deficiency, which could potentially influence the observed outcomes. Forthcoming work should target CXCL13 in specific cell types using conditional knockout mice. Second, we have not quantified all the leukocyte subsets, such as macrophages, monocytes, and neutrophils, which may also respond to CXCL13. Third, defining the exact cause of death in survival studies is challenging, as pathological events may occur in organs beyond the lungs, which were not examined. Fourth, our study focused exclusively on young male mice, whereas the importance of CXCL13 may vary with age, gender, and comorbidities in both mice and humans. Future studies should investigate CXCL13-mediated T and B cell immune responses in the draining lymph nodes and spleen to provide insights into systemic immunity and immune memory formation. While CXCL13 deficiency has been associated with impaired immune cell recruitment and tertiary lymphoid structure formation. Their deficiency may influence the secretion of other chemokines involved in immune responses, and therefore, whether the CXCL13 deficiency is directly related to severe SARS-CoV-2 infection remains uncertain.

In conclusion, our study proposes that an early stimulation of the CXCL13/CXCR5 chemokine axis promotes CXCR5^+^CD19^+^ B-cell and follicular/effector CXCR5^+^CD4^+^ T-cell responses in the lungs associated with protection from severe and fatal COVID-19 following Infection with the pathogenic SARS-CoV-2 Delta variant. While the underlying mechanism of the CXCL13/CXCR5 chemokine axis-mediated protection in SARS-CoV-2 infection remains to be fully elucidated, the report offers a potential new immunotherapeutic target for treatment.

## ACKNOWLEDGEMENTS

The authors would like to thank UC Irvine’s Medical Center and Institute for Clinical and Translational Science (ICTS) for helping with blood drawing from COVID-symptomatic and asymptomatic individuals.

